# phyC: Clustering cancer evolutionary trees

**DOI:** 10.1101/069302

**Authors:** Yusuke Matsui, Atsushi Niida, Ryutaro Uchi, Koshi Mimori, Satoru Miyano, Teppei Shimamura

## Abstract

**Motivation:** Multi-regional sequencing provides new opportunities to investigate genetic heterogeneity within or between common tumors from an evolutionary perspective. Several state-of-the-art methods have been proposed for reconstructing cancer sub-clonal evolutionary trees based on multi-regional sequencing data to develop models of cancer evolution. However, the methods developed thus far are not sufficient to characterize and interpret the diversity of cancer sub-clonal evolutionary trees.

**Results:** We propose a clustering method (`phyC`) for cancer sub-clonal evolutionary trees, in which sub-groups of the trees are identified based on topology and edge length attributes. For interpretation, we also propose a method for evaluating the diversity of trees in the clusters, which provides insight into the acceleration of sub-clonal expansion. Simulation showed that the proposed method can detect true clusters with sufficient accuracy. Application of the method to actual multi-regional sequencing data of clear cell renal carcinoma and non-small cell lung cancer allowed for the detection of clusters related to cancer type or phenotype.

**Availability:** `phyC` is implemented with R(>=3.2.2) and is available from https://github.com/ymatts/phyC.

## 1 Introduction

Cancer is a heterogeneous disease. The genetic diversity is driven by several evolutionary processes such as somatic mutation, genetic drift, migration, and natural selection. The clonal theory of cancer (Nowell, 1976) is based on Darwinian models of natural selection in which genetically unstable cells acquire a somatic single nucleotide variant (SSNV), and selective pressure results in tumors with a biological fitness advantage for survival.

The development of multi-regional sequencing techniques has provided new perspectives of genetic heterogeneity within or between common tumors (Schuh *et al*, 2012; Newurger *et al.*, 2013; Carter *et al.*, 2012; Campbell *et al.*, 2008; Yachida *et al.*, 2010). The read counts from multi-region tumor and matched normal tissue sequences from each patient are then used to infer the tumor composition and evolutionary structure from variant allele frequencies (VAFs); *i.e.*, the proportion of variant alleles that contains SSNVs. Using the VAF, the cancer evolutionary histories can be reconstructed as a tree, termed a sub-clonal evolutionary tree, which reflects how the identified SSNVs are accumulated for each patient.

Recently, new state-of-the-art methods have been proposed for reconstructing cancer sub-clonal evolutionary trees based on multi-regional sequencing data to develop models of cancer evolution, and have been applied to many types of cancers in multiple patients (Jiao *et al.*, 2013; Roth *et al.*, 2014; Miller *et al.*, 2014; Zare *et al.*, 2014;, Strino *et al.*, 2013; Qiao *et al.*, 2014; Salem *et al.*, 2015; Popic *et al.*, 2015). Mostofthe reconstruction methods developed to date are based on two assumptions: (i) no mutation occurs twice in the course of cancer evolution, and (ii) no mutation can be lost (Nik-Zainal *et al.*, 2012) (Figure 1A). In this tree, the root and its subsequent node represent a normal cell and founder cell, respectively. Sub-clones are described as nodes below the founder cell, and edge lengths indicate the number of SSNVs that are newly accumulated in descendant nodes (Figure 1B). For details, see Beerenwinkel *et al.* (2014).

**Fig 1.**
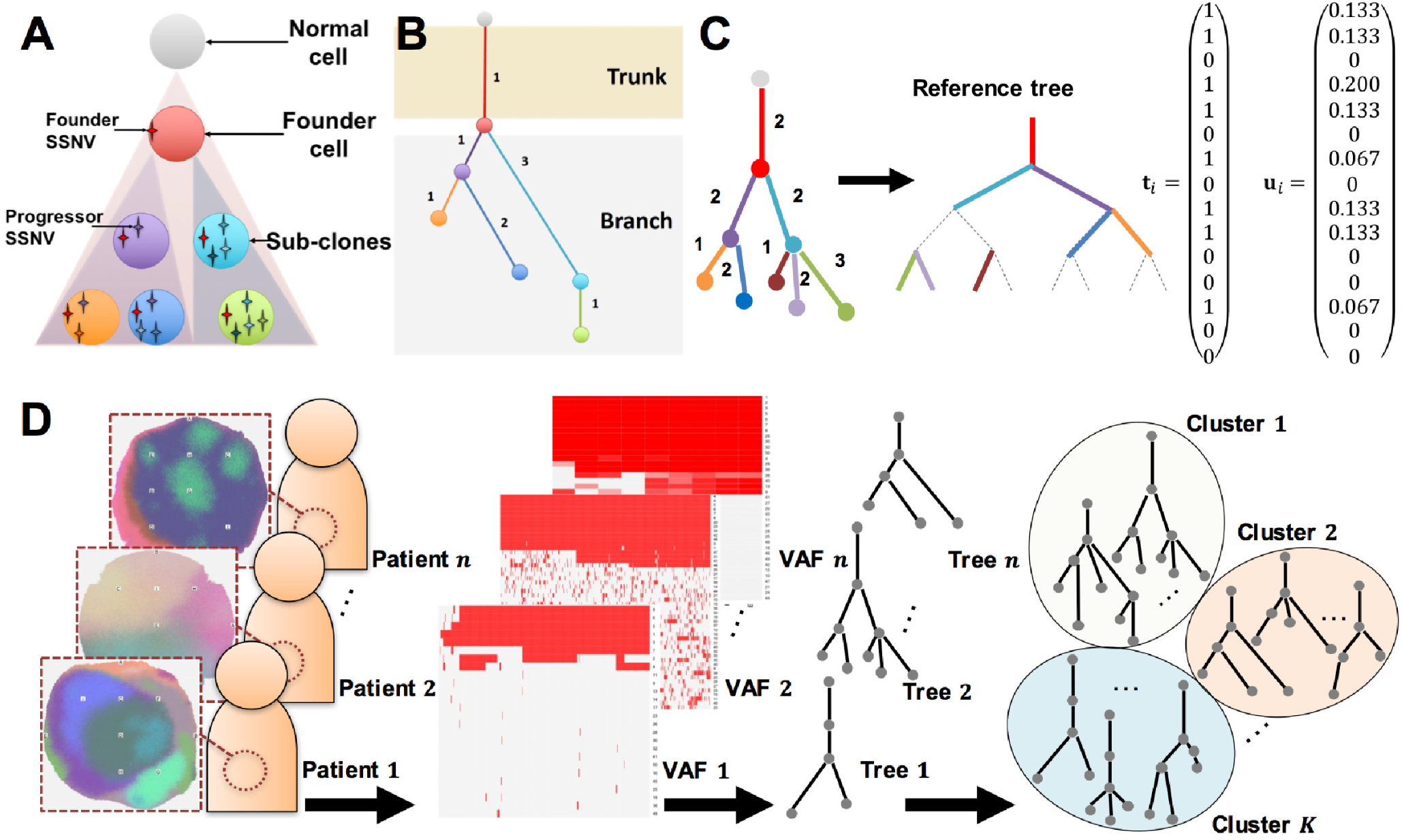
Overview of the proposed method. **(A)** Example of sub-clonal evolution. A founder cell is established after a normal cell acquires several passenger mutations and driver mutations (founder SSNVs), and sub-clones evolve by acquiring progressor SSNVs. Each color (purple, orange, dark blue, light blue, and green) of circles represents different sub-clones. **(B)** Example of sub-clonal evolutionary tree in the case of (A). A root and immediate node represent the normal cell and founder cell, respectively. Subsequent nodes indicate sub-clones and edge lengths indicate the number of SSNVs acquired in sub-clones. **(C)** Example of the registration of a tree. To resolve (p1)-(p4) for comparison of the evolutionary trees, a sufficiently large bifurcated tree is constructed, which is the reference tree (note that we have omitted bifurcation from the root for clearer visualization). The tree topologies and attributes are mapped to the reference tree beginning with those with largest depth to those with the smallest depth. In the case of a tie, the sub-trees are mapped from those with the largest edge length. Zero-length edges are regarded as degenerated edges (dashed lines). Edge lengths are normalized by the sum of all edge lengths within tumors. The resulting trees can be represented as topology vectors **t**_*i*_ and edge length vectors **u**_*i*_. (**D**) Clustering cancer evolutionary trees to summarize the evolutionary history of cancer for each patient. The trees are reconstructed based on the VAFs and then *n* cancer sub-clonal evolutionary trees are divided into *K* subgroups based on tree topologies and edge attributes. Through the registration, *n* evolutionary trees can be represented as *m*-dimensional *n* vectors in Euclidean space, and a standard clustering algorithm can be applied.

Although the reconstruction methods developed thus far have revealed intra-tumor heterogeneity by analysis of individual sub-clonal evolutionary trees, methods for obtaining a detailed understanding of inter-tumor heterogeneity according to evolutionary patterns with a set of the trees are lacking. A variety of sub-clonal evolutionary trees can be considered as the consequence of underlying tumor evolutionary principle, which may lead to resistance to chemotherapeutics and targeted therapies (Swanton, 2014; Venkatesan and Swanton, 2016). Characterizing inter-tumor heterogeneity by patterns of evolutionary tree is important for developing new targeted therapies and for preventing the emergence of drug resistance. Several studies have suggested specific evolutionary patterns of tumors with various and at times conflicting result. For example, Gerlinger *et al.* (2014) identified parallel evolution of sub-clones in clear cell renal cell carcinomas (ccRCCs), whereas no such parallel evolution was evident in studies on non-small cell lung cancer (NSCLC) (de Bruin *et al.*, 2014; Zhang *et al.*, 2014). Zhang *et al.* (2014) also showed that in a relapsed group of patients, the fraction of SSNVs in sub-clones was significantly larger than that of founder cells. These studies indicate that both the sub-clonal branching patterns and fraction of SSNVs in sub-clones are important factors for identifying the cancer type or phenotype of related subgroups.

In this paper, we propose a new clustering method for cancer sub-clonal evolutionary trees based on tree topologies and edge attributes that describe the relationships of sub-clones and the number of SSNVs that accumulate in the sub-clones (Figure 1D). Our conceptual framework is based on object-oriented data analysis (Marron, *et al.*, 2014), in which the observation units are non-numeric objects such as functions and trees.

Comparison of phylogenetic trees has long been discussed in the context of the evolution of species, and several comparative analytical methods have been developed, including nearest neighbor interchanging (Waterman and Smith, 1978), subtree transfer distance (Allen and Steel, 2001), quartet distance (Estabrook *et al.*, 1985), Robinson-Foulds distance (Robinson and Foulds, 1981), path length metrics (Steel and Penny, 1993), and Billera-Holmes-Vogtmann (BHV) distance (Billera, *et al.*, 2001). However, these distances are defined for phylogenetic trees with the same set of leaves, and therefore cannot accurately deal with the following problems, (p1)-(p4), that are specific to the context of cancer sub-clonal evolutionary trees.

**(p1)** The parental sub-clone has one child sub-clone or more than two child sub-clones (no bifurcation)
**(p2)** The number of sub-clones varies among patients (different numbers of leaves)
**(p3)** SSNV contents differ among patients (different leaf labels)
**(p4)** The number of detected SSNVs varies among patients (bias of edge attributes)

These problems motivated us to develop a method for transforming tree objects via transformation of the tree topologies and edge attributes to allow for effective comparison among trees, a procedure we refer to as tree registration. Given the complexity of cancer evolution, strict comparison of observed cancer sub-clonal evolutionary trees is unrealistic. In particular, the structures and sizes of observed evolutionary trees differ owing to the wide variation in sub-clonal evolution, and because the subclones themselves contain numerous progressor SSNVs that differ among patients. Alternatively, we focus on the biologically important features that may be related to cancer type- or phenotype-associated evolutionary structures, such as drug-sensitive sub-clonal evolution. Moreover, caner progression can be considered as the consequence of a complex sequence of several evolutionary events such as somatic mutation, genetic drift, migration, and natural selection, which do not happen at the same rate for every patient. Accordingly, for comparison of cancer evolutionary trees between two patients, the relative number of SSNVs should be considered rather than the fixed number of SSNVs.

The main contributions of this paper are development of (i) a tree registration method for cancer evolutionary trees, (ii) a clustering method of the registered trees, and (iii) an evaluation method of the clusters, which can be applied using our software `phyC` in the **R** environment.

In the registration, we resolve the issues raised in (p1)–(p4) though development of a method for transforming tree objects by mapping tree topologies and their attributes to make the trees comparable (figure 1C). The registered trees are embedded in Euclidean space, which enables defining the distance between the cancer sub-clonal evolutionary trees. Based on this distance, we divide a set of the trees into several subgroups with a clustering method (figure 1D). We developed two tools for interpretation of the clusters: multidimensional scaling (MDS) and a sub-clonal diversity plot.

We evaluated the performance of `phyC` using simulated VAFs that mimic the process of cancer sub-clonal evolution under the framework of cellular automaton (Niida, *et al.*, 2015). We also demonstrate the applicability of `phyC` using two actual datasets to show the interpretability of the clustering results: a ccRCC (Gerlinger *et al.*, 2014) and NSCLC (Zhang *et al.*, 2014) dataset.

`phyC` is implemented with R(>= 3.2.2) and is available from https://github.com/ymatts/PhyC.

## 2 Methods

We denote *n* reconstructed cancer sub-clonal evolutionary trees as *X* = {*x_i_*; *i* = 1, 2,…, *n*}, and the edges and edge lengths are denoted as {*e_ij_*; *i* = 1, 2,…, *n*, *j* = 1, 2,…, *m_i_*} and {|*e_ij_* |; *i* = 1, 2,…,*n*}, respectively. Without loss of generality, {*e*_*i*_ı; *i* = 1, 2,…, *n*} indicates the edge from the normal cell to the founder cell. Given the number of terminal nodes *N*_*i*_; *i* = 1, 2,…,*n*, we define the shortest path that is a set of edges connecting the root and terminal nodes with the minimum number of edges, denoted as *p_ik_* ={{*e*_*il*_; *i* = 1, 2,…, *n*,*l* ∈ *P_ik_* }; *k* = 1, 2,…,*N_i_*}, where *P*_*ik*_ ⊆ {1, 2,…, *m*_*i*_}. We define *depth* as the number of edges in the shortest path, and we denote *d_ik_* = *n*(*P*_*ik*_), where *n*(·) represents the number of elements in a set.

### 2.1 Registration

We developed a registration method for cancer evolutionary trees. The goal of the registration is to transform the observed trees such that dissimilarities can be defined with consideration of the tree topologies and edge attributes. To solve the problems (p1)–(p4), we provide the following approaches, (q1) and (q2):

**(q1)** Reference tree encoding
**(q2)** Normalizing edge lengths

To account for (p1)–(p3), we consider a reference tree-encoding approach. In this approach, we prepare a very large bifurcated tree called a reference tree (corresponding to as the maximum tree in Feragen *et al.*, 2013) and encode the observed tree topologies and edge lengths onto the reference. Zero-length edges are regarded as degenerated edges (Figure 1C). The advantage of this approach is that once we encode the observed tree onto a reference tree, the comparison can be simply achieved for trees of the same structures and sizes. To account for the issue (p4), we developed a method for normalization of the edge length to remove the bias of the detected number of SSNVs.

Here, we describe the details of the registration method. First, we set the maximum depth in *X* as *d_max_*(*X*) =max {*d_ik_*; *i* = 1, 2,…, *n*, *k* = 1, 2,…,*N_i_*} and define the reference tree as follows.

Definition 1 (Reference tree). *The reference tree is a bifurcated tree with the minimum depth of d_max_*(*X*).

Thus, the reference tree has *m* = 2(2^*d_max_*(*X*)^ – 1) edges (figure 1C). We denote the reference tree as *X_ref_* with edges and edge lengths *E_k_*; *k* = 1, 2,…,*m* and |*E_k_*|;*k* = 1, 2,…,*m*, respectively. The registration can then be defined with the reference tree.

Definition 2 (Registration). *Registration is a mapping f: X* ↦ *X_ref_*.

We define the mapped trees as *Y* = {*f* (*x_i_*); *i* = 1, 2,…, *n*}, and more specifically, the mapped edge and edge length are set as {*E_ik_*; *i* = 1, 2,…, *n*, *k* = 1, 2,…, *m*} and {|*E_ik_*|; *i* = 1, 2,…, *n*, *k* = 1, 2,…, *m*}, respectively. The number of edges differs between the observed tree and the reference tree, and we also need to account for any unmapped edges. Since the degenerated edges can be regarded as the zero-length edge when considering the distance of trees (Feragen *et al.*,2013), we can define |*E*_*ik′*_|= |*e_ij_*|; *k*′ ∈ *A* for the mapped edge index set *A* ⊆ {1, 2,…, *m*} and | *E_ik′_*| = 0; *k*′ ∈ *B* for the unmapped edge index *B* = {1, 2,…, *m*}\*A*.

To resolve (p3), we developed the mapping rule *eĳ* ↦ *E_ik_* for *j* = 1, 2,…, *m_i_, k* = 1, 2,…, *m*, such that the observed trees are mapped onto the reference tree begining with sub-trees with the largest depths to those with the smallest depths (Figure 1C). When the depths are the same among the sub-trees, we use the edge length and map the sub-trees beginning with the largest edge lengths.

In the last step of the registration, we perform normalization for the edge length. Zhang *et al.* (2014) importantly suggested that the ratio of the number of SSNVs in a trunk to that in the sub-clones is related to certain phenotypes such as whether or not the cancer is recurrent. Therefore, we consider that the ratio of the number of accumulated SSNVs is an important factor to characterize and compare the cancer sub-clonal evolutionary trees, and we divided each edge length by the total number of SSNVs within patients.

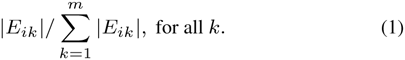

### 2.2 Clustering set of registered trees

To define the dissimilarity between the registered trees, we begin with the space of the set of the registered trees. Theoretically, the space of a registered trees is the special case of BHV space (Birella, *et al.*, 2001), where the space of a semi-labeled tree is a disjointed Euclidean space. In BHV space, each tree topology corresponds to a Euclidean orthant, and if the tree topologies are the same, then they will lie in the same orthant where the coordinates correspond to the edge lengths. Since we only encode the observed trees onto the reference tree with the same topology, the registered trees do indeed lie in the same orthant.

Corollary 1 (Euclidean embedding). *The registered trees lie in Euclidean space.*

We represent the registered tree *y_i_* as a vector in Euclidean space as follows. The binary vector is defined as 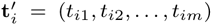; *t_ij_* ∈ {0, 1}, whose elements correspond to the presence of each edge. If an edge is present, then *t*_*ij*_ = 1; otherwise, *t*_*ij*_ = 0. We also define edge attributes, whose elements correspond to each edge length as 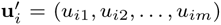; *u_ij_* ∈ **R**. Using **t** and **u**, the tree **y**_*i*_ can be represented as

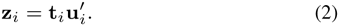

We set 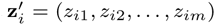; *z_im_*∈ **R**. Thus, *n* registered trees are represented as the *n* × *m* matrix 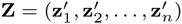.

We define the dissimilarity as follows:

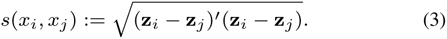

The basic statistics of the cancer sub-clonal evolutionary trees can also be defined. The tree average is defined as 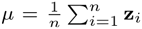 and the tree variance is defined as 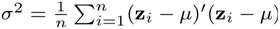.

Based on the tree representation in (2), which can be regarded as *n* observations with an *m* features matrix, we can simply apply standard clustering algorithms and divide the *n* trees into subgroups. Hierarchical clustering was then implemented using `phyC`.

### 2.3 Graphical representation

Interpreting clustering results is a key issue for tree comparison, which requires understanding the features of the cancer sub-clonal evolutionary trees in clusters. In particular, visual representation can be a powerful tool for such interpretation. Therefore we developed two computational tools for comparing trees and understanding the cluster features.

#### MDS

To effectively compare the trees, we approximately embedded the registered trees into lower-dimensional Euclidean space. For this purpose, we applied classical MDS (CMDS) (Torgerson, 1952), which is a dimension-reduction technique based on singular value decomposition. We omit the details of the CMDS algorithm and briefly describe the method below. Given the symmetric distance matrix *S* = {*s_ij_*; *i, j* =1,2,…, *n*}, the double centered matrix

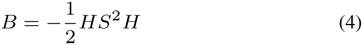

is positive semi-definite where 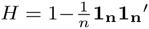, and can be diagonalized as

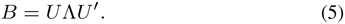

The constructed coordinates are obtained by 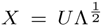. For this purpose, we use the distance that is defined in (3). CMDS requires knowing the number of dimensions, which we set to two for the purpose of convenient visualization. In `phyC`, we overlaid the tree shapes over the coordinates and visually compared the tree structures based on dissimilarity.

#### Sub-clonal diversity plot

To visualize how sub-clones evolve with respect to SSNV accumulation, we apply the concept of a lineage-through-time (LTT) plot, which is commonly used for visualizing the timing of speciation events in studies of the birth-death process. The LTT plot generally describes the time vs. number of lineages; in the present case, this is expressed as the number of sub-clones (y-axis) vs. the fraction of accumulated SSNVs (x-axis), and the plot is referred to as a sub-clonal diversity plot. In the plot, *y* = 0 means that there is no sub-clone, and thus only a normal cell exists, and *y* = 1 indicates that there is a founder cell. For example, (*x, y*) = (0.3, 1) indicates that the founder cell is established with the accumulation of 30% SSNVs. For *y* > 1, the growth curve in the plot represents how many sub-clones emerged for a given fraction of SSNVs. If the curve is upright, the sub-clones evolve with a small fraction of SSNV accumulation, and conversely, if the curve grows with gradual steps, the sub-clones acquire a relatively large fraction of SSNVs. A gradual growth curve was observed in the case of parallel evolution shown in Gerlinger *et al.*, (2014), which will be demonstrated below in the implementation of the ccRCC dataset.

## 3 Results

### 3.1 Simulation data

We evaluated the performance of the proposed method using simulation data. We generated a VAF profile of multi-region sequences using the BEP simulator (Niida *et al.*, 2015; Uchi *et al.*, 2016) based on a cellular automaton model. Three main parameters were used to mimic cancer sub-clonal evolution: mutation rate (*m*), the number of driver genes (*d*), and strength of the driver mutation gene (*f*). The simulator starts with one normal cell and grows in a probabilistic manner. In Niida *et al.* (2015), the mutation rate was identified as one of the most important factors contributing to cancer heterogeneity.

We assume two clusters consisting of 30 cancer sub-clonal evolutionary trees. Using the BEP simulator, we generated tumor VAF profiles over nine regions. The evolutionary patterns are controlled with the mutation rate m, and following Niida *et al.* (2015), we set *m* = 0.01 (parameter setting 1) and *m* = 0.0001 (parameter setting 2) for each cluster, which represents the cancer heterogeneity characterized as neutral evolution and Darwinian evolution, respectively. Other parameters were set as *d* = 8 and *f* = 1.2 in both clusters. The evolutionary trees were constructed using Wagner parsimony trees from the VAF profiles with the function `acctran` in the **R** package `phangorn` (Klaus, 2011), after converting the VAF profiles to binary profiles: if *VAF* ≥ 0.05, then we regarded the gene as mutated and set it to 1; otherwise, it was considered to be not mutated and was set to set 0.

We performed the registration for 60 trees and calculated the dissimilarities among the trees. Figure 2 shows the MDS plot based on the dissimilarity, and each color represents the parameter settings 1 (light green) and 2 (purple). From the MDS plot, the generated evolutionary trees appeared to consist of the two clusters.

**Fig 2.**
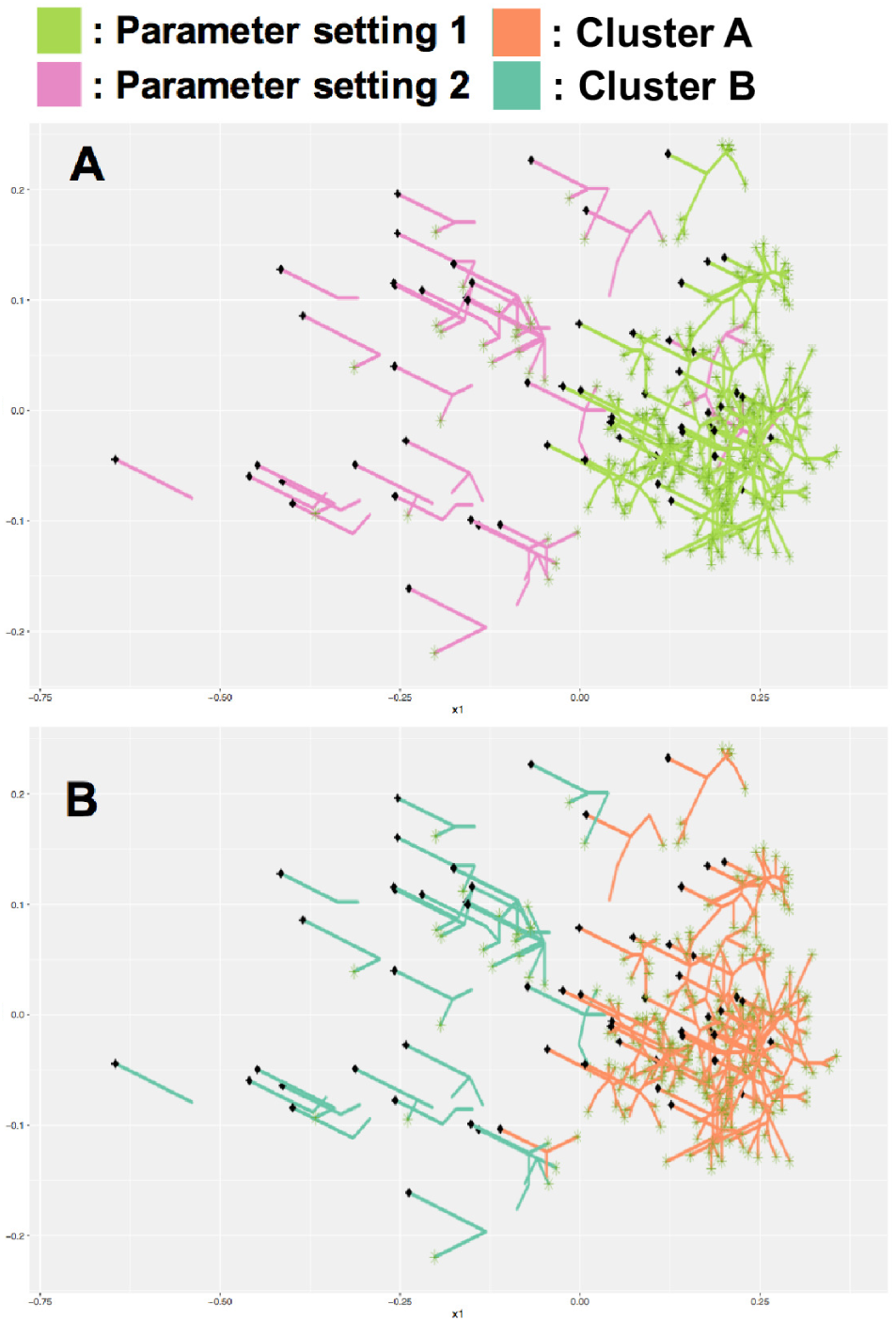
**(A)** True clusters and evolutionary trees generated from the BEP simulator. Trees in parameter setting 1 and 2 are grouped according to high and low mutation rates. **(B)** Clusters detected by `phyC`. Trees with each parameter were effectively classified into true clusters, except for four trees. The true positive rate was 93%.

We performed hierarchical clustering with Ward’s method (Ward, 1963) on the 60 registered trees, which were divided into two subgroups: A and B. Although four trees of parameter setting 2 were misclassified into cluster A, our method could effectively classify the sub-clonal evolutionary trees with a true positive rate of 93% (Table 1).

**Table 1.** Clustering result of simulation data

|  | Parameter setting 1 | Parameter setting 2 |
| --- | --- | --- |
| Cluster A | 30 | 4 |
| Cluster B | 0 | 26 |

The simulation result demonstrated that `phyC` could accurately classify sub-clonal structures derived from different evolutionary principles, based on their evolutionary tree shapes.

### 3.2 Real data

We here demonstrate the application of our proposed method using an actual ccRCC dataset (Gerlinger *et al.*, 2014) and NSCLC dataset (Zhang *et al.*, 2014), consisting of 8 and 11 multi-regional tumor samples and their VAFs were collected among 587 and 7,026 SSNVs, respectively. Since both studies used the maximum parsimony method to reconstruct the cancer sub-clonal evolutionary trees, we also adopted this method to analyze the datasets with our approach. We binarized the VAF profiles with *VAF* ≥ 0.05 as one and otherwise zero. Using the binary profile, we estimated phylogenetic trees using the function `acctran` in the **R** package phangorn and we obtained 19 cancer sub-clonal evolutionary trees.

#### ccRCC dataset

The ccRCC dataset consists of 8 sub-clonal evolutionary trees with clinical information related to treatments. We divided the 8 sub-clonal evolutionary trees of the ccRCC dataset into 3 subgroups using hierarchical clustering with Ward’s method. Table 2 shows the clustering result and a configuration of the 8 trees with CMDS (Figure 3A-1).

**Fig 3.**
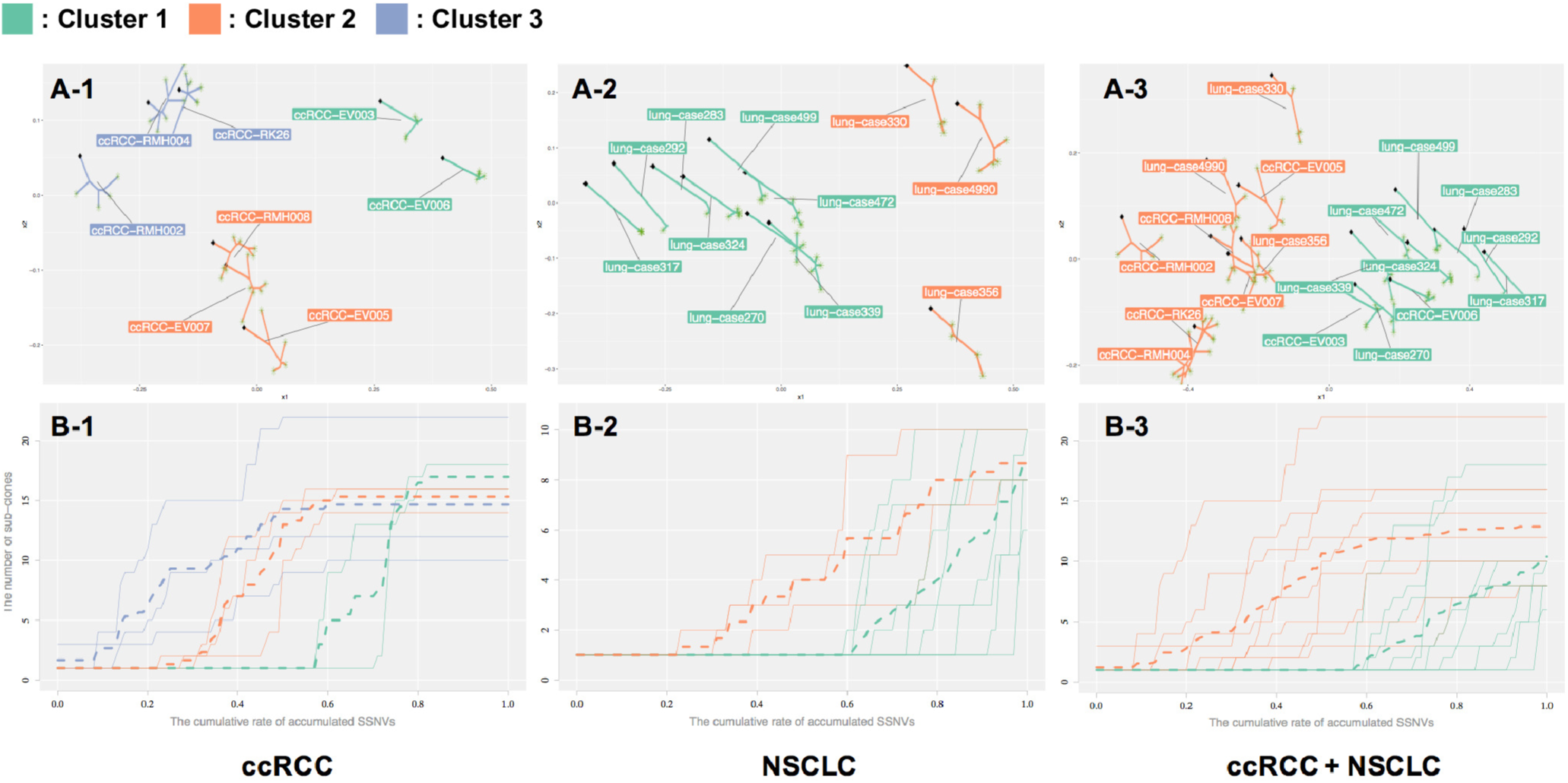
**(A-1)** Three clusters of the ccRCC dataset. Clusters 1 (green) and 2 (orange) reflect drug-sensitive sub-clonal evolution and parallel evolution, respectively; we cannot provide valid interpretation for cluster 3 (purple) at present. **(B-1)** Sub-clonal diversity plot of the ccRCC dataset. Each color corresponds to the clusters shown in (A). Sub-clonal expansions in cluster 1 occurred with *x* = 0.6; (*i.e.*, the proportion of SSNVs in the trunk is 60%). This result is in contrast to that obtained for cluster 2 (*x* = 0.2) and cluster 3 (*x* = 0.1). Trees in cluster 2 show gradual growth of diversity curves, indicating that these sub-clones acquire a relatively large fraction of SSNVs. The sub-clones independently evolve in spatially distinct regions (Gerlinger *et al.*, 2014. **(A-2)** Two clusters in the NSCLC dataset. Clusters 1 (green) and 2 (orange) reflect the non-recurrent and recurrent group, respectively. Only case 270 and case 356 were misclassified to clusters 1 and 2, respectively. **(B-2)** Sub-clonal diversity plot of the NSCLC dataset. Each color corresponds to the clusters shown in (A-2). **(A-3)** Two clusters in the ccRCC and NSCLC datasets combined. Clusters 1 (green) and 2 (orange) represent the cancer types NSCLC and ccRCC, respectively. **(B-3)** Sub-clonal diversity plot of the ccRCC and NSCLC datasets.

**Table 2.** Clustering result of the ccRCC dataset

| Cluster | Sample name |
| --- | --- |
| Cluster 1 | EV003, EV006 |
| Cluster 2 | EV005, EV007, RMH008 |
| Cluster 3 | RK26, RMH004, RMH002 |

Cases EV003 and EV006 in cluster 1 received pretreatment with everolimus (Gerlinger *et al.* 2014). The sub-clonal diversity plot shown in Figure 3B-1 demonstrates that sub-clonal expansions in cluster 1 occurred after 60% of the SSNVs accumulated, in contrast to cluster 2 (20%) and cluster 3 (10%), which may indicate that the drug interrupts acquisition of SSNVs in the sub-clones, leading to lower genetic diversity of the subclones. Cluster 1 reflects the drug-sensitive sub-clones group, and the result corresponds to the interpretation provided by Gerlinger *et al.* (2014).

Cases EV005, EV007, and RMH008 in cluster 2 acquired a large fraction of SSNVs in the sub-clones (Figure 3B-1). Comparison of the original tree shape demonstrated that the long branches are followed by several private branches. These three samples were reported as subclones of parallel evolution, *i.e.*, each sub-clone independently evolved in spatially distinct regions (Gerlinger *et al.*, 2014). Cluster 3, including RK26, RMH004, and RMH002, acquired the largest fraction of SSNVs among sub-clones; however, there is no valid interpretation for this result.

These findings demonstrate that our method can produce interpretable clusters for the drug-sensitive group and parallel evolution group in the ccRCC data set.

#### NSCLC dataset

The NSCLC dataset consists of 11 sub-clonal evolutionary trees with the following clinical information: staging (IA, IIA, IIIA, and IB), smoking status (former, current, and never), and recurrence (yes and no). We divided the 11 trees into 2 subgroups using hierarchical clustering with Ward’s method. The clustering result is shown in Table 3, and the configuration of CMDS is shown in Figure 3A-2.

**Table 3.** Clustering result of the NSCLC dataset

| Cluster | Sample name |
| --- | --- |
| Cluster 1 | case 317, case 292, case 283, case 324, case 499<br>case 472, case 339, case 270 |
| Cluster 2 | case 330, case 4990, case 356 |

Case 330 and case 4990 in cluster 2 are labeled as the recurrent group, but case 356 is not. As shown in Figure 3B-2, there are several long horizontal regions in the diversity curve, which indicates that each subclone acquired a large portion of SSNVs. This implies that the sub-clones contained a large fraction of the SSNVs after diverging from the founder cell, *i.e.*, representing a genetically new generation. Zhang *et al.* (2014) reported a similar observation, in which they found a significant difference (t-test) in the average fractions of SSNVs between the recurrent group and non-recurrent group. Case 356 is labeled as non-recurrent; however, it shows a large fraction of SSNVs in the sub-clones compared to that of the non-recurrent group, leading to a similar tree shape to that of trees of the recurrent group.

Cluster 1 consists of non-recurrent cases, except for case 270. Figure 3B-2 shows small horizontal regions in the diversity curve, which indicates that each sub-clone acquired a smaller portion of SSNVs compared to that observed in the sub-clones of cluster 2; that is, most of the SSNV acquisitions events had already occurred as founder mutations. Although case 270 is labeled as recurrent, it shows a lower fraction of SSNVs in the sub-clones compared to that of the recurrent group, and as a result, its tree shape resembles that of trees of the non-recurrent group.

The difference between the two clusters indicates that acquisition of a large fraction of SSNVs in sub-clones may influence the survival of cancer patients, and this feature could be captured and classified according to analysis of tree shapes through the proposed method.

#### Comparison of the ccRCC and NSCLC datasets

In addition to establishing the evolutionary pattern of a certain cancer type or sub-type, it is also interesting to compare the sub-clonal evolutionary patterns of different types of cancer. Therefore, we compare the sub-clonal evolutionary trees derived from the ccRCC and NSCLC datasets. We applied `phyC` to the 19 trees reconstructed as described in the previous sub-sections, which were divided into two distinct clusters (Table 4, Figure 3A-3).

**Table 4.** Clustering result of the ccRCC and the NSCLC datasets

| Cluster | Sample name |
| --- | --- |
| Cluster 1 | case 317, case 292, case 283, case 324, case 499<br>case 472, case 339, case 270, EV003, EV006 |
| Cluster 2 | EV005, EV007, RK26, RMH002, RMH004<br>RMH008, case 330, case 4990, case 356 |

The trees could be mainly classified according to cancer type. One of the main features of the ccRCC sub-clonal evolutionary trees in cluster 2 was the acquisition of a large fraction of SSNVs in the sub-clones, leading to the tendency of parallel evolution. In contrast to the ccRCC trees, the NSCLC sub-clonal evolutionary trees showed that a large fraction of SSNVs was acquired in the trunk, and not in the sub-clones, which confirmed that the important event had already occurred in the early stage of SSNVs acquisition (Zhang *et al.*, 2014).

Some of the trees were classified with different cancer types, including case 330, case 4990, and case 356 in cluster 2, and EV003 and EV006 in cluster 1. Case 330 and case 4990 in the NSCLC dataset are part of the recurrent group, and the tree shape of case 356 is similar to that of the recurrent group (Table 4). Furthermore, their sub-clonal evolutionary trees were similar to the ccRCC trees, which implies that new aberrant SSNVs related to recurrence, which were not present during early SSNVs accumulation, were acquired in sub-clones. EV003 and EV006 in the ccRCC dataset are samples of drug-sensitive tumors, and their tree shapes resemble those of NSCLC trees, which further supports that the drug interrupts the accumulation of SSNVs in the sub-clones.

## 4 Discussion

Considering the generally high level of inter-tumor heterogeneity, it is important to be able to identify phenotype- or cancer type-related sub-clonal evolutionary patterns. Previous studies have classified and interpreted the branching patterns of such sub-clones with manual methods, and then separately analyzed the compositions of SSNVs in each sub-clone. However, development of a quantitative analysis method is required to best scale datasets containing a large number of samples with sub-clonal evolutionary trees for characterizing and interpreting the evolutionary patterns.

Our approach relies on reconstruction methods of sub-clonal evolutionary trees, and we used a parsimony approach that is widely adopted in studies of multi-regional sequencing. The proposed method only requires knowledge of the edges and edge attributes of rooted trees, and is therefore widely applicable to outputs of other recently developed state-of-the-art reconstruction methods, which allowed us to consider the heterogeneous mixture of cells within a sample.

There are several limitations of the present method that are worth mentioning, which should be tackled in further investigations. First, we have ignored the specific content of SSNVs in the sub-clones. We believe that this is a reasonable assumption to some extent, since the variation of SSNVs is too large to yield an effective comparison. However, the effects and consequences of different types of SSNVs can also vary, such as driver mutations or passenger mutations. Thus, when comparing edges with the same lengths from different trees, the two edges may not actually be equivalent if driver genes are included in one edge but not in the other. Therefore, a method that can incorporate the effect of driver genes in the sub-clones should be explored in future work.

Second, we have here considered only the SSNVs accumulated into sub-clonal evolutionary trees, ignoring potential copy number or epigenetic aberrations; however, these factors may also affect heterogeneity within a tumor. Multi-regional sequence analysis has been performed using exome sequencing as well as copy number, methylation, and mRNA expression array profiling, providing an integrated interpretation of cancer sub-clonal evolution (Uchi *et al.*, 2016). To determine the sub-clonal evolutionary patterns from these integrated data, our method can be extended to the case of multivariate edge attributes, including copy number variations and hyper- or hypo-methylation, as well as other genetic and epigenetic aberrations.

Finally, we did not take into account the potential effects of regional sampling biases and individual variations among tumors or patients. Gerlinger *et al.* (2014) pointed out that increasing the sequenced regions of samples might lead to additional detection of sub-clones, and thus the complexity of inferred sub-clonal evolutionary trees might be affected by the sampling strategy. Therefore, a method for sampling bias reduction is needed to improve the clustering accuracy and plausible interpretation.

## 5 Conclusion

We developed `phyC`, which was designed for clustering a set of cancer sub-clonal evolutionary trees to characterize cancer sub-clonal evolutionary patterns according to tree shape, based on analysis of tree topologies and edge attributes. Using this approach, we effectively identified the evolutionary patterns with different degrees of heterogeneity in a simulation study. We also successfully detected the phenotype-related and cancer type-related subgroups when applying this method to actual ccRCC and NSCLC data. Our method represents the first practical method to quantitatively and accurately compare a variety of sub-clonal evolutionary trees with different structures, sizes, and labels, and with biases of edge length, while further allowing for biological interpretation. Our results imply that this approach has potential applications for personalized medicine such as predicting outcomes of chemotherapeutics and targeted therapies, *e.g.*, drug-resistance, based on sub-clonal evolutionary trees and we believe that the value and impact of our work will grow as more and more multi-regional sequencing datasets of patients become available.

